# Metabolic state modulates neural processing of odors in the human olfactory bulb

**DOI:** 10.1101/2023.08.23.554431

**Authors:** Behzad Iravani, Johannes Frasnelli, Artin Arshamian, Johan N. Lundström

## Abstract

The olfactory and endocrine systems have recently been shown to reciprocally shape the homeostatic processes of energy intake. As demonstrated in animal models, the individual’s metabolic state dynamically modulates how the olfactory bulb process odor stimuli using a range of endocrine signals. Here we aimed to determine whether the neural processing of odors in human olfactory bulb is modulated by metabolic state. Participants were exposed to food-associated odors, in separate sessions being hungry and sated, while neural responses from the olfactory bulb was obtained using electrobulbogram. We found significantly higher gamma power activity (51-100Hz) in the OB’s response to odors during the Hunger compared to Sated condition. Specifically, EBG gamma power were elevated while hungry already at 100ms after odor onset, thereby suggesting intra-bulbar modulation according to metabolic state. These results demonstrate that, akin to other animal models, hunger state affects OB activity in humans. Moreover, we show that the EBG method has the potential to measure internal metabolic states and, as such, could be used to study specificities in olfactory processing of individuals suffering from pathologies such as obesity or anorexia.

## Introduction

One of the primary functions of the olfactory sense is the detection and identification of food sources (Stevenson, 2010). Consequently, it is expected that the feeding state (i.e., the metabolic state), ranging from hunger to satiation, has an influence on olfactory function. For instance, fasting rats exhibit an enhanced capacity for olfactory detection compared to sated rats (Aime et al., 2007). Feeding state is also reflected at the cerebral level. Specifically, the olfactory bulb (OB) □— the first processing stage in the central olfactory pathway □— demonstrates feeding state-dependent responses. In hungry rats, exposure to a food odor elicits a hyperpolarization in the mitral cell layer compared to sated states which mostly elicits a depolarization (Pager et al., 1972; Royet & Pager, 1981). Consequently, the output from the olfactory bulb is highly dependent on feeding state: mitral cell output as measured in the olfactory peduncle and the lateral olfactory tract is maximal in hungry animals and minimal in sated animals (Pager et al., 1972; Royet & Pager, 1981). Although it is well established that the olfactory bulb demonstrates metabolic state-dependent responses in animal models, whether metabolic state influence neural processing in the human olfactory bulb is not known.

Akin to animal models, human participants demonstrate heightened olfactory sensitivity when experiencing hunger compared to satiation (Stafford & Welbeck, 2011). Specifically, fasting participants exhibited greater sensitivity in olfactory testing, particularly within the peri-threshold range (Cameron et al., 2012; Hanci & Altun, 2016; Ramaekers et al., 2016; Albrecht et al., 2009). This indicates that human olfactory performance as a function of feeding state mirrors behavior observed in animal models and lends support to the notion that the neural signal in the human OB is modulated by the individual’s metabolic status. To date, this question has yet to be explored, primarily due to the absence of non-invasive methods to assess human OB function. The Electrobulbogram (EBG), our recently developed method, measures human OB function via an electrophysiological signal recorded with 4 active electroencephalography electrodes placed on the forehead of individuals (Iravani et al., 2020). The measurement is a reliable and valid sign of OB processing of odors in both young and older adults (Iravani, Arshamian, Schaefer, et al., 2021). Crucially, the EBG detects an early evoked gamma synchronization around 100ms in response to odors, the cortical source of which is localized in the OB (Iravani et al., 2020). Importantly, we have in multiple experiments excluded other signal sources than the OB (Iravani et al., 2020). In rodents, such gamma oscillations are linked to processes within the OB (Wilson & Sullivan, 2011). The EBG can hence be used to map the communication between the OB and higher order olfactory processing structures (Iravani, Arshamian, Lundqvist, et al., 2021).

Here we use the EBG method to investigate the electrophysiological difference in the human OB between hungry and sated states. We hypothesized that the feeding state modulates gamma power of the OB signal in response to olfactory stimulation.

## Materials and Methods

### Participants

A total of 17 individuals (mean age: 30 ± SD 5 years, 10 women) participated in this study. All participants declared themselves as generally healthy, of a healthy weight, non-smokers with no history of head trauma (associated with unconsciousness) and indicated to not suffer from any present or past neurological, nutritional, or metabolic disorders. Participants BMI ranged between 17-27 with a mean of 23.1 (SD 2.9); 4 participants were classified as overweight. Prior to inclusion, all participants provided written informed consent and we further determined that all participants had a normal sense of smell in using a 16-item 4-alternative forced-choice odor identification task (Landis et al., 2004). All participants identified at least 13 out of the 16 odors. All aspects of the study were approved by the University of Pennsylvania’s Institutional Review Board (IRB approval #832080) and the protocol complies with the revised Declaration of Helsinki.

### Metabolic state manipulation

Participants participated in two testing sessions, being either hungry or sated, with two days in-between. In both sessions, they arrived at the lab at 9am without having eaten or drinking anything other than water since 6pm the evening before. In the *Hungry* condition, participants initiated the study and underwent odor EEG testing. All participants reported that they followed the dietary restriction. In the *Sated* condition, participants were presented with a variety of energy bars, none containing the two odors, and asked to eat as many as they wanted until they felt sated, **Figure 1A**. On average, participants ate 2.5 bars (range 1-4). In the Sated condition, odor testing did not start until 30 minutes after participants reported themselves as sated to allow time for the metabolic state to initiate its potential influence on the olfactory system. Starting order in respect of conditions was decided for each participant in a counter-balanced order based on enrollment while ensuring even sex distribution between conditions.

**Figure 1.**
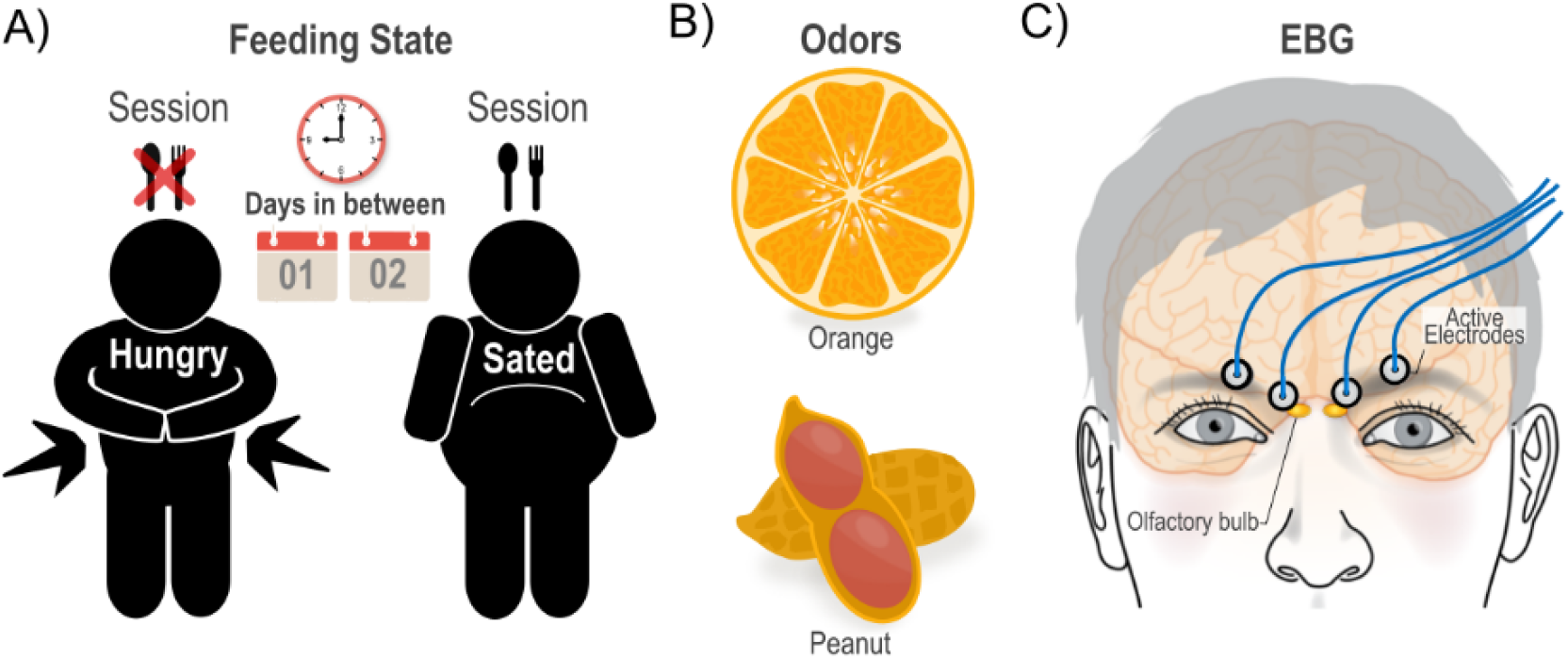
Experimental design, odor stimuli, and EBG. **A**) Summary of the experimental design illustrating the two recording sessions with two different degrees of feeding states for each participant. **B**) Odor stimuli (Orange and Peanut). **C**) The signal from 4 active EBG electrodes is recorded during odor smelling task. The electrodes are placed on the forehead, slightly above the eyes, as it is shown here, following the outline of the person’s eyebrows.

### Odor stimuli and odor delivering methods

We selected two food related odors, namely *Orange* Oil (Sigma Aldrich, # W282510, CAS 8008-57-9) and *Peanut* Oil (Takasago Natural and Artificial Peanut Flavor, TAK-053887), **Figure 1B**. We diluted the *Orange* and *Peanut* odors in odorless silica-filtered, light mineral oil (CAS 8042-47-5) to a volume/volume concentration of 30% and 35%, respectively. These dilutions were rated as iso-intense in a pilot experiment (n=15; *t*(14) = 0.29, *p* = .77, *CI* -0.58 0.43). Odors were presented using a computer-controlled olfactometer with a rise-time (i.e., the time needed for the odorant to reach 90% of the maximum concentration) of 200ms (Lundstrom et al., 2010). Odors were delivered bilaterally by switching odorized air (flow rate: 1.5 L/min per side) into a constant stream of clean air (flow rate: 0.5 L/min; rendering a total flow per nostril of 2 L/min) to minimize somatosensory co-stimulation due to the odor onset. To limit potential effects of respiration, participants were instructed to breathe through their mouth during the whole time of the experiment and odors were delivered passively and non-synchronized to breathing cycle. Participants’ task was to rate odor intensity after each stimulation throughout experiment. However, since the behavioral data for intensity ratings of 7 participants were corrupted, we could not use this variable for further analysis.

All the timing, odor triggering, and data logging were implemented in E-prime 2 (Psychology Software Tools, Pennsylvania). The trials’ onset was jittered (800–2000ms) before the odor presentation with a duration of 1000ms to prevent participants from predicting odor onset. Moreover, to minimize the effect of habituation we kept a random inter-trial interval of 9500 – 19500ms.

### Electrobulbogram recording and preprocessing

The electrophysiological signal was sampled at 512Hz using 32 EEG scalp electrodes (ActiveTwo, BioSemi, Amsterdam, The Netherland), placed according to international 10/20 standard, an additional 4 active EBG electrodes (Iravani et al., 2020) placed on the forehead, **Figure 1C**, and 3 electrodes to record eye movement and blinks. Data was band-pass filtered at 0.01–100Hz during recording within the ActiView software (BioSemi, Amsterdam, The Netherland). Before recording, we ascertained that the offset of electrodes was below 40mV by visually inspecting the electrodes’ offset and adjusting those with a high offset to meet our criterion.

The pre-processing (MATLAB 2019b) of the EBG signals started with epoching data from 100ms pre-stimulus to 1500ms post-stimulus. We further re-referenced data from the mandatory ActiveTwo systems reference to mastoid electrodes followed by notch filtered the EBG data at 50Hz to remove the line noise and its two harmonics. Moreover, the data were band-pass filtered to 1-100Hz using 4thorder Butterworth filter to remove any drift or non-electrophysiological activity. After this initial pre-processing step, we removed trials with either muscle or blink artifacts using ocular inspection of each trial. Hence, for determining whether a trail contains muscle artifact or not, we first filtered the data to 110-140 Hz using 8th order Butterworth filter, followed by Hilbert transform to extract the instantaneous amplitudes. Subsequently, the amplitudes were z-scored and the trials that surpassed the threshold of 8 were identified as muscle artifacts. Similarly, detecting blinks was implemented by filtering the signal to 1-15 Hz using 4th order Butterworth filter, Hilbert transform and z-scoring. We identify trails with blink artifact if the z-scored amplitude of filtered signal surpassed the threshold of 4 and trials containing either muscle or blink artifact were excluded from further analysis.

### Time-frequency analysis of Electrobulbogram data

Artifact-less EBG trials were then decomposed into time-frequency components using multi-taper-convolution method implemented in field trip toolbox 2018 (Oostenveld et al., 2011), within MATLAB 2019b. Specifically, we estimated the power of gamma band (i.e., 30-100Hz) with step of 0.5Hz and in total for 141 frequency bins. Subsequently, the time interval of 100ms pre-stimulus to 250ms post-stimulus was selected as the time window of interest and we explored this interval of interest with a step size of 5ms. We used a discrete prolate spheroidal sequence as for the window function and set the smoothing parameters to 50% of a given frequency which yield in using of two windows for each frequency bin. We further removed trials that had a power spectral density with 3 standard deviations (absolute values) larger than average. Next, trials were averaged for each participant separately, followed by scale-based normalization (Randolph, 2006) and converting to decibels. The width of the window was selected such that it covered the whole trial length.

### Statistical analysis

To test the effect of energy level on the OB power spectrum we averaged the two odors’ spectrogram and used a non-parametric Monte Carlo permutation test to test the significance between the two conditions *Hungry* and *Sated*. A 5000-permutation test was performed with a prior significance detection criterion of *p* < .05 and K > 100, where K denotes the number of significant bins in the cluster. Type 1 error was controlled by cluster correction. We further assessed the generalizability of our findings using a post-hoc Jackknife resampling test. Accordingly, the power values from the significant cluster were extracted and contrasted against two conditions. In each iteration of the Jackknife test one participant was left out and the average for remaining subject was computed. After all subjects were once left out, the power value of the iterations was averaged, and 95% CI was estimated.

### Data availability

The data underlying this article cannot be shared publicly due to the ethical approval and participant consent the data was acquired under. The data will, however, be shared on reasonable request to the corresponding author.

## Results

We addressed the main question, whether feeding state modulated olfactory bulb activity, by contrasting OB gamma band power spectrum in the two conditions (Hunger > Sated). We focused on the gamma band because past studies have connected gamma oscillations in human OB to odor processing (Iravani, Arshamian, Lundqvist, et al., 2021; Iravani et al., 2020; Iravani, Arshamian, Schaefer, et al., 2021) and, in the animal model literature, gamma processing has been linked to within area processing and afferent ‘bottom-up’ communication (Frederick et al., 2016; van Kerkoerle et al., 2014). We found higher gamma power in the OB during the Hunger condition compared to Sated. Specifically, EBG gamma power were elevated around 51-100Hz already at 100ms after odor onset (**Figure 2A**). We further assessed whether the increase in gamma power was a significant effect using 5000-resampling Monte Carlo tests with significant detection criteria of p < .05, K >100. We found that the increase of gamma power for Hungry compared to Sated state was statistically significant (*t* = 2.36, *p* < .006, *p*-value confidence range: [.005 .007], **Figure 2B**).

**Figure 2.**
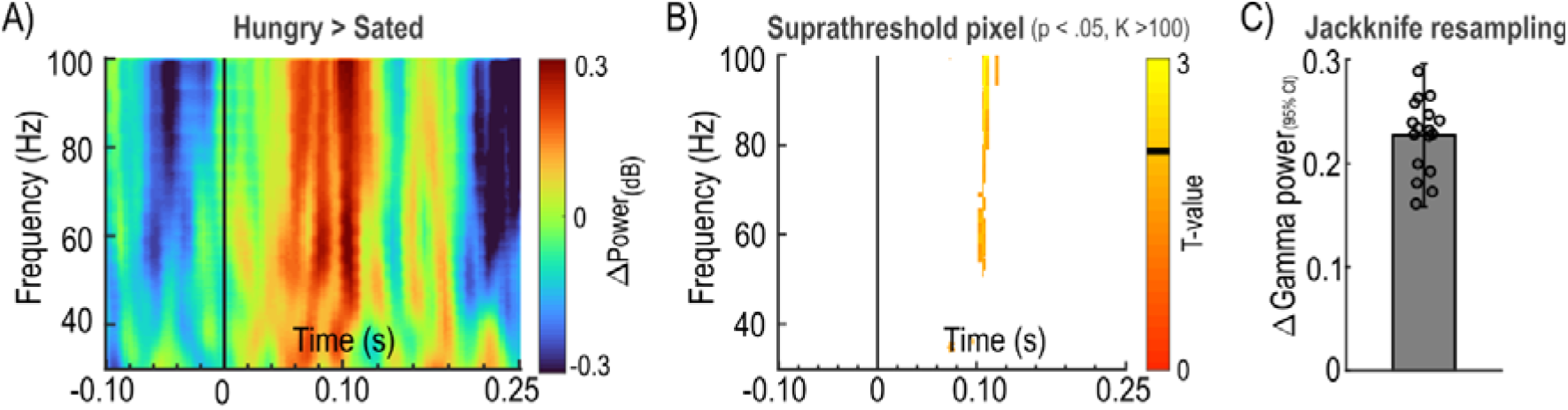
OB gamma power and hunger state. **A**) The spectrogram of the EBG channels demonstrated more power in gamma band at around 100ms post odor onset for “Hungry” session compared with “Sated” session. The vertical black line indicates the onset of odor, and the change of power values are transformed to decibels and color-coded. The warmer colors show higher power for “Hungry” whereas the cooler colors show higher power for “Sated”. **B**) The t-map from 5000 Monte Carlo resampling indicates a significant cluster for “Hungry” compared to “Sated” in the time-frequency bins around 51-100Hz and 100ms. Warmer colors indicate higher t-values with black line in color bar indicating significance threshold. **C**) The mean power of significant cluster was resampled using Jackknife method to estimate the 95% CI. The bar shows the mean over the resampled iterations. The circles represent the mean value for a given iteration and the error bar shows the Jackknife 95% CI.

Finally, to assess the stability of our obtained results, we also assessed the variation in our obtained gamma power results using Jackknife resampling to ascertain that the result is not driven by any specific combination of individuals included in this study. Here, the resampled averaged power values for the significant cluster were all above 0 (resampled Jackknife 95% CI: [0.16, 0.30],**Figure 2C**). Hence, the grand average and its 95% CI over the Jackknife iterations were deemed as unbiased effect of energy level on the OB gamma power values.

## Discussion

Here we show that the human OB, like other mammals, encodes the individual’s feeding status. Specifically, we show an early and significantly higher gamma power in the EBG signal when participants were hungry compared to when they were sated. As early gamma power in OB mainly reflects processing of olfactory stimulus, with minimal top-down contribution from other central nervous structures, this suggests that the observed increase in gamma power from sated to hungry also mainly reflects processing within the OB (Frederick et al., 2016; Martin & Ravel, 2014). Importantly, early gamma activity has also been associated with odor valence perception in humans (Iravani, Schaefer, et al., 2021). Our results suggest that the human olfactory bulb is governed by similar mechanisms as in other animals, such as rats. In hungry rats, exposure to a food odor significantly increases activity in the mitral cell layer compared to sated rats (Pager et al., 1972; Royet & Pager, 1981).

Feeding state modulates behavior and perception of olfactory stimuli. Fasted rats are more sensitive to avoid the odor they were negatively conditioned with than sated individuals (Aime et al., 2007). Fasted rats also explore toys odorized with food odors significantly longer than sated ones (Prud’homme et al., 2009). Likewise, humans that are hungry perform better in a variety of olfactory tasks (Cameron et al., 2012; Hanci & Altun, 2016; Ramaekers et al., 2016; Stafford & Welbeck, 2011), although food odors may be less well detected when hungry (Albrecht et al., 2009; Stafford & Welbeck, 2011). Along the same lines, feeding state also influences pleasantness of odors, as sated individuals rate food odors as less pleasant (Albrecht et al., 2009; Small et al., 2001). If these changes in odor perception are a result of activity within the OB alone, or induced by top-down piriform projections is unknown. However, studies have shown that the secondary olfactory cortex, such as the orbitofrontal cortex, change its neural processing of odors from hunger to satiation (Small et al., 2001). Moreover, It should be noted that while gamma oscillations from the olfactory bulb induce gamma activation upstream in the piriform cortex (Osinski et al., 2018), where they for example contribute to the perception of odor identity (Iravani, Arshamian, Lundqvist, et al., 2021; Yang et al., 2022), the piriform cortex in turn projects top-down information about odor identity to the OB via delta and theta oscillations (Iravani, Arshamian, Lundqvist, et al., 2021). Thus, many internal processes within the olfactory network can contribute to the control of metabolic states. Nonetheless, we argue that our early gamma activity in the OB mainly reflects bottom-down information. For this to happen, the OB must be able to code metabolic state directly. Different mechanisms have been put forward on how feeding state influences olfactory processing.

Several molecules related to feeding state and behavior act on the olfactory system and, crucially, the OB (Palouzier-Paulignan et al., 2012). Most studied among feeding-related molecules are the orexigenic molecules ghrelin (Kojima et al., 1999) and orexins A/B (Sakurai et al., 1998), as well as the anorexigenic molecule insulin. Ghrelin is produced in the gastrointestinal tract and is a hormone that triggers hunger and initiates feeding behavior (Palouzier-Paulignan et al., 2012). Indeed, ghrelin receptors can be found in different layers of the OB and, accordingly, ghrelin enhances olfactory sensitivity in both rodents and humans (Tong et al., 2011). Likewise, food odors are perceived as more pleasant after ghrelin injection (Han et al., 2018). Orexins A and B are neuropeptides in hypothalamic neurons that are involved in different neuroendocrine functions, including food intake (Palouzier-Paulignan et al., 2012). These neurons also project to the OB (Hardy et al., 2005). Exposure to orexin A alters firing rate and frequency of OB neurons (Apelbaum et al., 2005; Hardy et al., 2005; Prud’homme et al., 2009) and increases olfactory sensitivity (Julliard et al., 2007). On the other side of the spectrum, the anorexigenic hormone insulin, has its highest cerebral concentration in the OB (Baskin et al., 1983) where insulin receptors are widespread (Palouzier-Paulignan et al., 2012). Intravenous injection of insulin increases insulin levels in the OB in rats to levels observed in satiation and abolishes the effect of hunger on olfactory detection thresholds (Aime et al., 2012; Kuczewski et al., 2014). Other hormones and endogenous modulators that potentially modulate olfactory bulb activity in hunger and satiation include orexigenic neuropeptide Y, endocannabinoids, endogenous opioids as well as anorexigenic leptin, cholecystokinin, and others (see (Palouzier-Paulignan et al., 2012) for an overview). Our results, and this body of literature, is in line with the notion of olfaction playing a central role in flavor processing (Auvray & Spence, 2008) and therefore in regulating eating behavior (Yeomans, 2000). It is, however, not clear to what extent any of these molecules are involved in the electrophysiological modulation of OB activity we have observed here. However, that they are involved in OB processing has been established and might serve as a potential mechanism for feeding-state modulation of the human OB. Future studies should seek to manipulate the concentration of these peptides in the body and measure its effect on the OB activity directly.

Past studies have demonstrated a rich link between metabolic disorders and olfactory processing. Among others, it has been demonstrated that olfactory impairment is prevalent among individuals with obesity (Campolo et al., 2021) and that BMI modulates neural processing in both cortical and subcortical (Jacobson et al., 2019) neural areas. Although our obtained results cannot be directly extended to individuals with metabolic disorders, the clear link between metabolic state and odor processing in the olfactory bulb suggest that the area is implicated in the mechanism of how metabolic disorders alters odor perception. Assessing this link is ripe for future investigations.

Our study has specific limitations: while we asked participants to evaluate odors with regards to intensity and pleasantness, data from nearly half of the participants (7 out of 17) perceptual ratings were digitally corrupted which prevented us to investigate potential interesting links between odor perception, metabolic state, and olfactory bulb activity. Along the same line, the study was setup to assess our research question in a dichotomized manner and only statement of satiety or hunger was collected. Assessing whether odor-dependent OB responses linearly scale with either biomarkers or subjective ratings of hunger/satiety, although partly outside the scope of the preset study and requiring a much larger sample, would provide a deeper mechanistic understanding of how OB processing of odors are shaped by feeding states. Second, only two odors were used meaning that we could not assess to what extent caloric content associated with the food odor modulates the effect. Although a separate question, this is the natural next step. Third, although employing a within participant design, a relatively small number of participants were included into our experiment. Nevertheless, by performing non-parametric permutation tests, we were able to control for false positives. Together with the Jackknife resampling approach, this increased the generalizability of our results and demonstrated stable data. Future studies aiming to replicate our results should include a larger sample size, a wider variety of odors associated with food of different caloric content, assess influence of feeding state on odor intensity and pleasantness, as well as subsequent OB processing.

In conclusion, we show that hunger state affects OB activity in humans, as assessed by the EBG. From a methodological perspective our finding has implications for researchers using the EBG as higher signal-to-noise ratio may be obtained in hungry participants. In the broader aspect, we show that our method has the potential to measure internal metabolic states and as such it could be used to study the specificities in olfactory processing of individuals suffering from pathologies such as obesity or anorexia.

## Acknowledgment

Funding provided by the Knut and Alice Wallenberg Foundation (KAW 2018.0152), awarded to JNL. JF is supported by CIHR, NSERC, and FRQS.

## AI declaration

No AI tool, or function, was used when analyzing the data or when composing the manuscript.

## References

Aime, P., Duchamp-Viret, P., Chaput, M. A., Savigner, A., Mahfouz, M., & Julliard, A. K. (2007). Fasting increases and satiation decreases olfactory detection for a neutral odor in rats. Behav Brain Res, 179(2), 258–264. 10.1016/j.bbr.2007.02.012

Aime, P., Hegoburu, C., Jaillard, T., Degletagne, C., Garcia, S., Messaoudi, B., Thevenet, M., Lorsignol, A., Duchamp, C., Mouly, A. M., & Julliard, A. K. (2012). A physiological increase of insulin in the olfactory bulb decreases detection of a learned aversive odor and abolishes food odor-induced sniffing behavior in rats. PLoS ONE, 7(12), e51227. 10.1371/journal.pone.0051227

Albrecht, J., Schreder, T., Kleemann, A. M., Schopf, V., Kopietz, R., Anzinger, A., Demmel, M., Linn, J., Kettenmann, B., & Wiesmann, M. (2009). Olfactory detection thresholds and pleasantness of a food-related and a non-food odour in hunger and satiety. Rhinology, 47(2), 160–165. https://www.ncbi.nlm.nih.gov/pubmed/19593973

Apelbaum, A. F., Perrut, A., & Chaput, M. (2005). Orexin A effects on the olfactory bulb spontaneous activity and odor responsiveness in freely breathing rats. Regul Pept, 129(1-3), 49–61. 10.1016/j.regpep.2005.01.003

Auvray, M., & Spence, C. (2008). The multisensory perception of flavor. Conscious Cogn, 17(3), 1016–1031. 10.1016/j.concog.2007.06.005

Baskin, D. G., Porte, D., Jr., Guest, K., & Dorsa, D. M. (1983). Regional concentrations of insulin in the rat brain. Endocrinology, 112(3), 898–903. 10.1210/endo-112-3-898

Cameron, J. D., Goldfield, G. S., & Doucet, E. (2012). Fasting for 24 h improves nasal chemosensory performance and food palatability in a related manner. Appetite, 58(3), 978–981. 10.1016/j.appet.2012.02.050

Campolo, J., Corradi, E., Rizzardi, A., Parolini, M., Dellanoce, C., Di Guglielmo, M. L., Tarlarini, P., Cattaneo, M., Trivella, M. G., & De Maria, R. (2021). Correlates of olfactory impairment in middleaged non-diabetic Caucasian subjects with stage I-II obesity. European Archives of Oto-Rhino-Laryngology, 278(6), 2047–2054. 10.1007/s00405-020-06442-5

Frederick, D. E., Brown, A., Brim, E., Mehta, N., Vujovic, M., & Kay, L. M. (2016). Gamma and Beta Oscillations Define a Sequence of Neurocognitive Modes Present in Odor Processing. J Neurosci, 36(29), 7750–7767. 10.1523/JNEUROSCI.0569-16.2016

Han, J. E., Frasnelli, J., Zeighami, Y., Larcher, K., Boyle, J., McConnell, T., Malik, S., Jones-Gotman, M., & Dagher, A. (2018). Ghrelin Enhances Food Odor Conditioning in Healthy Humans: An fMRI Study. Cell Rep, 25(10), 2643–2652 e2644. 10.1016/j.celrep.2018.11.026

Hanci, D., & Altun, H. (2016). Hunger state affects both olfactory abilities and gustatory sensitivity. Eur Arch Otorhinolaryngol, 273(7), 1637–1641. 10.1007/s00405-015-3589-6

Hardy, A. B., Aioun, J., Baly, C., Julliard, K. A., Caillol, M., Salesse, R., & Duchamp-Viret, P. (2005). Orexin A modulates mitral cell activity in the rat olfactory bulb: patch-clamp study on slices and immunocytochemical localization of orexin receptors. Endocrinology, 146(9), 4042–4053. 10.1210/en.2005-0020

Iravani, B., Arshamian, A., Lundqvist, M., Kay, L. M., Wilson, D. A., & Lundstrom, J. N. (2021). Odor identity can be extracted from the reciprocal connectivity between olfactory bulb and piriform cortex in humans. Neuroimage, 237, 118130. 10.1016/j.neuroimage.2021.118130

Iravani, B., Arshamian, A., Ohla, K., Wilson, D. A., & Lundstrom, J. N. (2020). Non-invasive recording from the human olfactory bulb. Nat Commun, 11(1), 648. 10.1038/s41467-020-14520-9

Iravani, B., Arshamian, A., Schaefer, M., Svenningsson, P., & Lundstrom, J. N. (2021). A non-invasive olfactory bulb measure dissociates Parkinson’s patients from healthy controls and discloses disease duration. NPJ Parkinsons Dis, 7(1), 75. 10.1038/s41531-021-00220-8

Iravani, B., Schaefer, M., Wilson, D. A., Arshamian, A., & Lundstrom, J. N. (2021). The human olfactory bulb processes odor valence representation and cues motor avoidance behavior. Proc Natl Acad Sci U S A, 118(42). 10.1073/pnas.2101209118

Jacobson, A., Green, E., Haase, L., Szajer, J., & Murphy, C. (2019). Differential Effects of BMI on Brain Response to Odor in Olfactory, Reward and Memory Regions: Evidence from fMRI. Nutrients, 11(4). 10.3390/nu11040926

Julliard, A. K., Chaput, M. A., Apelbaum, A., Aime, P., Mahfouz, M., & Duchamp-Viret, P. (2007). Changes in rat olfactory detection performance induced by orexin and leptin mimicking fasting and satiation. Behav Brain Res, 183(2), 123–129. 10.1016/j.bbr.2007.05.033

Kojima, M., Hosoda, H., Date, Y., Nakazato, M., Matsuo, H., & Kangawa, K. (1999). Ghrelin is a growth-hormone-releasing acylated peptide from stomach. Nature, 402(6762), 656–660. 10.1038/45230

Kuczewski, N., Fourcaud-Trocme, N., Savigner, A., Thevenet, M., Aime, P., Garcia, S., Duchamp-Viret, P., & Palouzier-Paulignan, B. (2014). Insulin modulates network activity in olfactory bulb slices: impact on odour processing. J Physiol, 592(13), 2751–2769. 10.1113/jphysiol.2013.269639

Landis, B. N., Konnerth, C. G., & Hummel, T. (2004). A study on the frequency of olfactory dysfunction. Laryngoscope, 114(10), 1764–1769. 10.1097/00005537-200410000-00017

Lundstrom, J. N., Gordon, A. R., Alden, E. C., Boesveldt, S., & Albrecht, J. (2010). Methods for building an inexpensive computer-controlled olfactometer for temporally-precise experiments. Int J Psychophysiol, 78(2), 179–189. 10.1016/j.ijpsycho.2010.07.007

Martin, C., & Ravel, N. (2014). Beta and gamma oscillatory activities associated with olfactory memory tasks: different rhythms for different functional networks? Front Behav Neurosci, 8, 218. 10.3389/fnbeh.2014.00218

Oostenveld, R., Fries, P., Maris, E., & Schoffelen, J. M. (2011). FieldTrip: Open source software for advanced analysis of MEG, EEG, and invasive electrophysiological data. Comput Intell Neurosci, 2011, 156869. 10.1155/2011/156869

Osinski, B. L., Kim, A., Xiao, W., Mehta, N. M., & Kay, L. M. (2018). Pharmacological manipulation of the olfactory bulb modulates beta oscillations: testing model predictions. J Neurophysiol, 120(3), 1090–1106. 10.1152/jn.00090.2018

Pager, J., Giachetti, I., Holley, A., & Le Magnen, J. (1972). A selective control of olfactory bulb electrical activity in relation to food deprivation and satiety in rats. Physiology & Behavior, 9(4), 573–579. 10.1016/0031-9384(72)90014-5

Palouzier-Paulignan, B., Lacroix, M. C., Aime, P., Baly, C., Caillol, M., Congar, P., Julliard, A. K., Tucker, K., & Fadool, D. A. (2012). Olfaction under metabolic influences. Chem Senses, 37(9), 769–797. 10.1093/chemse/bjs059

Prud’homme, M. J., Lacroix, M. C., Badonnel, K., Gougis, S., Baly, C., Salesse, R., & Caillol, M. (2009). Nutritional status modulates behavioural and olfactory bulb Fos responses to isoamyl acetate or food odour in rats: roles of orexins and leptin. Neuroscience, 162(4), 1287–1298. 10.1016/j.neuroscience.2009.05.043

Ramaekers, M. G., Verhoef, A., Gort, G., Luning, P. A., & Boesveldt, S. (2016). Metabolic and Sensory Influences on Odor Sensitivity in Humans. Chem Senses, 41(2), 163–168. 10.1093/chemse/bjv068

Randolph, T. W. (2006). Scale-based normalization of spectral data. Cancer Biomarkers: Section A of Disease Markers, 2(3–4), 135–144. 10.3233/cbm-2006-23-405

Royet, J. P., & Pager, J. (1981). Olfactory bulb responsiveness to an aversive or novel food odor in the unrestrained rat. Brain Res Bull, 7(4), 375–378. 10.1016/0361-9230(81)90032-0

Sakurai, T., Amemiya, A., Ishii, M., Matsuzaki, I., Chemelli, R. M., Tanaka, H., Williams, S. C., Richardson, J. A., Kozlowski, G. P., Wilson, S., Arch, J. R., Buckingham, R. E., Haynes, A. C., Carr, S. A., Annan, R. S., McNulty, D. E., Liu, W. S., Terrett, J. A., Elshourbagy, N. A., Yanagisawa, M. (1998). Orexins and orexin receptors: a family of hypothalamic neuropeptides and G protein-coupled receptors that regulate feeding behavior. Cell, 92(4), 573–585. 10.1016/s0092-8674(00)80949-6

Small, D. M., Zatorre, R. J., Dagher, A., Evans, A. C., & Jones-Gotman, M. (2001). Changes in brain activity related to eating chocolate: From pleasure to aversion. Brain, 124, 1720–1733.

Stafford, L. D., & Welbeck, K. (2011). High hunger state increases olfactory sensitivity to neutral but not food odors. Chem Senses, 36(2), 189–198. 10.1093/chemse/bjq114

Stevenson, R. J. (2010). An initial evaluation of the functions of human olfaction. Chem Senses, 35(1), 3–20. 10.1093/chemse/bjp083

Tong, J., Mannea, E., Aime, P., Pfluger, P. T., Yi, C. X., Castaneda, T. R., Davis, H. W., Ren, X., Pixley, S., Benoit, S., Julliard, K., Woods, S. C., Horvath, T. L., Sleeman, M. M., D’Alessio, D., Obici, S., Frank, R., & Tschop, M. H. (2011). Ghrelin enhances olfactory sensitivity and exploratory sniffing in rodents and humans. J Neurosci, 31(15), 5841–5846. 10.1523/JNEUROSCI.5680-10.2011

van Kerkoerle, T., Self, M. W., Dagnino, B., Gariel-Mathis, M. A., Poort, J., van der Togt, C., & Roelfsema, P. R. (2014). Alpha and gamma oscillations characterize feedback and feedforward processing in monkey visual cortex. Proc Natl Acad Sci U S A, 111(40), 14332–14341. 10.1073/pnas.1402773111

Wilson, D. A., & Sullivan, R. M. (2011). Cortical processing of odor objects. Neuron, 72(4), 506–519. 10.1016/j.neuron.2011.10.027

Yang, Q., Zhou, G., Noto, T., Templer, J. W., Schuele, S. U., Rosenow, J. M., Lane, G., & Zelano, C. (2022). Smell-induced gamma oscillations in human olfactory cortex are required for accurate perception of odor identity. PLoS Biol, 20(1), e3001509. 10.1371/journal.pbio.3001509

Yeomans, M. R. (2000). Rating changes over the course of meals: what do they tell us about motivation to eat? Neurosci Biobehav Rev, 24(2), 249–259. 10.1016/s0149-7634(99)00078-0

